# Microsaccade direction reveals the variation in auditory selective attention processes

**DOI:** 10.1101/2024.08.07.606838

**Authors:** Shimpei Yamagishi, Shigeto Furukawa

**Affiliations:** NTT Communication Science Laboratories, Kanagawa, Japan; Shizuoka Graduate School of Public Health, Shizuoka, Japan; Shizuoka General Hospital, Shizuoka, Japan

## Abstract

Selective spatial attention plays a critical role in perception in the daily environment where multiple sensory stimuli exist. Even covertly directing attention to a specific location facilitates the brain’s information processing of stimuli at the attended location. Previous behavioral and neurophysiological studies have shown that microsaccades, tiny involuntary saccadic eye movements, reflect such a process in terms of visual space and can be a marker of spatial attention. However, it is unclear whether auditory spatial attention processes that are supposed to interact with visual attention processes influence microsaccades and vice versa. Here, we examine the relationship between microsaccade direction and auditory spatial attention during dichotic oddball sound detection tasks. The results showed that the microsaccade direction was generally biased contralateral to the ear to which the oddball sound was presented or that to which sustained auditory attention was directed. The post-oddball modulation of microsaccade direction was associated with the behavioral performance of the detection task. The results suggest that the inhibition of stimulus-directed microsaccade occurs to reduce erroneous orientation of ocular responses during selective detection tasks. We also found that the correlation between microsaccade direction and neural response to the tone originated from the auditory brainstem (frequency-following response: FFR). Overall, the present study suggests that microsaccades can be a marker of auditory spatial attention and that the auditory neural activity fluctuates over time with the states of attention and the oculomotor system, also involving the auditory subcortical processes.

**Significant statement:** Microsaccades, tiny involuntary saccadic eye movements, reflect covert visual attention and influence neural activity in the visual pathway depending on the attention states. However, we lack convincing evidence of whether and how microsaccades reflect auditory spatial attention and/or neural activity along the auditory pathway. Intriguingly, we showed that the microsaccade direction exhibited systematic stimulus-related change and correlated with auditory brainstem frequency-following response (FFR) during the dichotic selective attention task. These results suggest that microsaccades are associated with general spatial attention processes, not restricted to the visual domain, and can be a good tool for accessing fluctuating neural activity that may covary with the attention states.

## Introduction

Our brain selectively processes sensory input based on its relevance to ongoing cognitive tasks, and such selective processing, often collectively referred to as attention, modulates the neural information processes in sensory modalities (e.g., auditory cortex: Fritz et al., 2007; visual cortex: Gilbert & Sigman, 2007). Attention is a dynamic process, and its states vary spontaneously over time for both visual and auditory domains (Esterman et al., 2013; Van Den Brink et al., 2016; Terashima et al., 2021). An objective way to capture fluctuating attention would assist in the understanding of the neural system of attention and track time-varying attention.

Microsaccades (MSs), tiny and involuntary saccadic eye movements, serve as a good candidate for a “window” for evaluating the attentional state. In visual tasks, MS direction correlates with covert attention: central cues indicating the target direction elicit MS towards the cued location (Engbert & Kliegl, 2003; Hafed & Clark, 2002), while peripheral cues elicit an early MS towards and a late MS away from the cued location (Hafed et al., 2021; Hafed & Ignashchenkova, 2013; Laubrock et al., 2005). In a simple detection task of a peripheral visual target without cueing, the short (or long) manual reaction time is accompanied by MS toward (or away from) the target (Tian et al., 2016a). These temporal dynamics of MSs may reflect temporal shifts in attention states and are associated with relevant neural substrates (Hafed et al., 2021). MS bias also can persist over a relatively long period depending on sustained visual attention (Xue et al., 2020) and can occur even for memorized visual items (Liu et al., 2022). Neurophysiological evidence suggests that visual cortical neuron firing rates increase following MS towards attended visual stimuli (Lowet et al., 2018).

In the auditory domain, however, links between attention and MS have been explored far less in phenomenology and neural substrates. The present study attempts to address the following issues. (1) Although general tendencies of cue-oriented MS have been reported during an auditory cueing task (Rolfs et al., 2005), it is unknown whether the MS reflects moment-by-moment fluctuation of attention states during auditory tasks and to what extent the shift of MS direction is linked to behavioral states during auditory attention tasks. (2) In contrast to the visual domain, there is a lack of evidence on how MSs are associated with neural activity in the auditory pathway. Exploration in the auditory domain should provide insights into the significance of MSs in supra- or cross-modal attention processes, as well as the utility of MSs as a biomarker for tracking auditory attention.

To address the two points raised above, we conducted a dichotic selective detection task: participants were asked to attend to the left or right ear during the presentation of a prolonged sequence of standard sounds and were required to detect the oddball sounds presented to the attended side. To observe the fluctuation of attention states over time, we used a sound sequence with a relatively long duration (2 – 3 minutes) compared to the previous study that examined the transient effect of the auditory cue on MS (Rolfs et al., 2005). We examined auditory neuronal activities by focusing on the auditory brainstem frequency-following response (FFR). Recent studies including source-level analysis of EEG responses showed that attention affects FFR to continuous speech stimuli (Forte et al., 2017; Etard et al., 2019; Price and Bidelman, 2021), while other studies reported a large individual variation in the effect of attention on the FFR (Lehmann and Schönwiesner, 2014), or even found that attention had no significant effect (Galbraith and Kane, 1993; Varghese et al., 2015). We hypothesized that attention-related brainstem activities, if any, are susceptible to the spontaneous temporal variations of the attention states, which could explain previous contradictory findings. We expected that if an MS property reflects instantaneous attentional states, we might observe attention-related FFR associated with the MS property.

## Material and Methods

### Participants

Human adults with normal hearing participated in three experiments; Experiment 1: nineteen participants (three males) with ages ranging from 22 to 49 years (mean = 38.53); Experiment 2: twelve participants (five males) with ages ranging from 20 to 48 years (mean = 37.08); and Experiment 3: twenty participants (thirteen males) with ages ranging from 20 to 28 years (mean = 22.1). The experimental protocols were approved by the Research Ethics Committee of Nippon Telegraph and Telephone (NTT) Communication Science Laboratories. All listeners gave written informed consent before the experiment.

### Apparatus

The eye data were recorded with an Eyelink system (ER Research, Toronto, Canada) at a sampling rate of 1000 Hz. Visual stimuli were generated with Matlab (R2016b) and were presented on a monitor with a resolution of 1920 × 1080 pixels and a refresh rate of 60 Hz. Participants sat on a chair and put their heads on a chin rest. The distance between the chin rest and the center of the display was around 70 cm. Auditory stimuli were synthesized with Matlab at a sampling frequency of 44.1 kHz and were presented binaurally through ER-3A insert earphones (Etymotic Research, Elk Grove Village, IL).

We used the ActiveTwo system (Biosemi, Amsterdam, Netherlands) to record EEG signals, from which frequency-following responses (FFRs) were derived. The electrophysiological responses were recorded differentially between Cz and both ear lobes (A1 and A2) with Ag-Ag/Cl active electrodes and sampled at 16384 Hz. The ActiveTwo system replaces the ground electrodes used in conventional EEG systems with two separate electrodes; Common Mode Sense (CMS) and Driven Right Leg (DRL) electrodes. FFRs were measured only in Experiments 1 and 3 (see below).

### Stimuli and procedure

In a typical experimental session, the participant’s task was to detect oddball sounds embedded in a sequence of standard sounds (complex tones) with alternating fundamental frequency (F0), presented to left and right ears (Fig.1). Table 1 summarizes the conditions of the session tested in the experiments. All three experiments examined the Dich_TargetL and Dich_TargetR conditions (see below), and there were additional tasks (as necessary) depending on the experiment.

**Figure 1.**
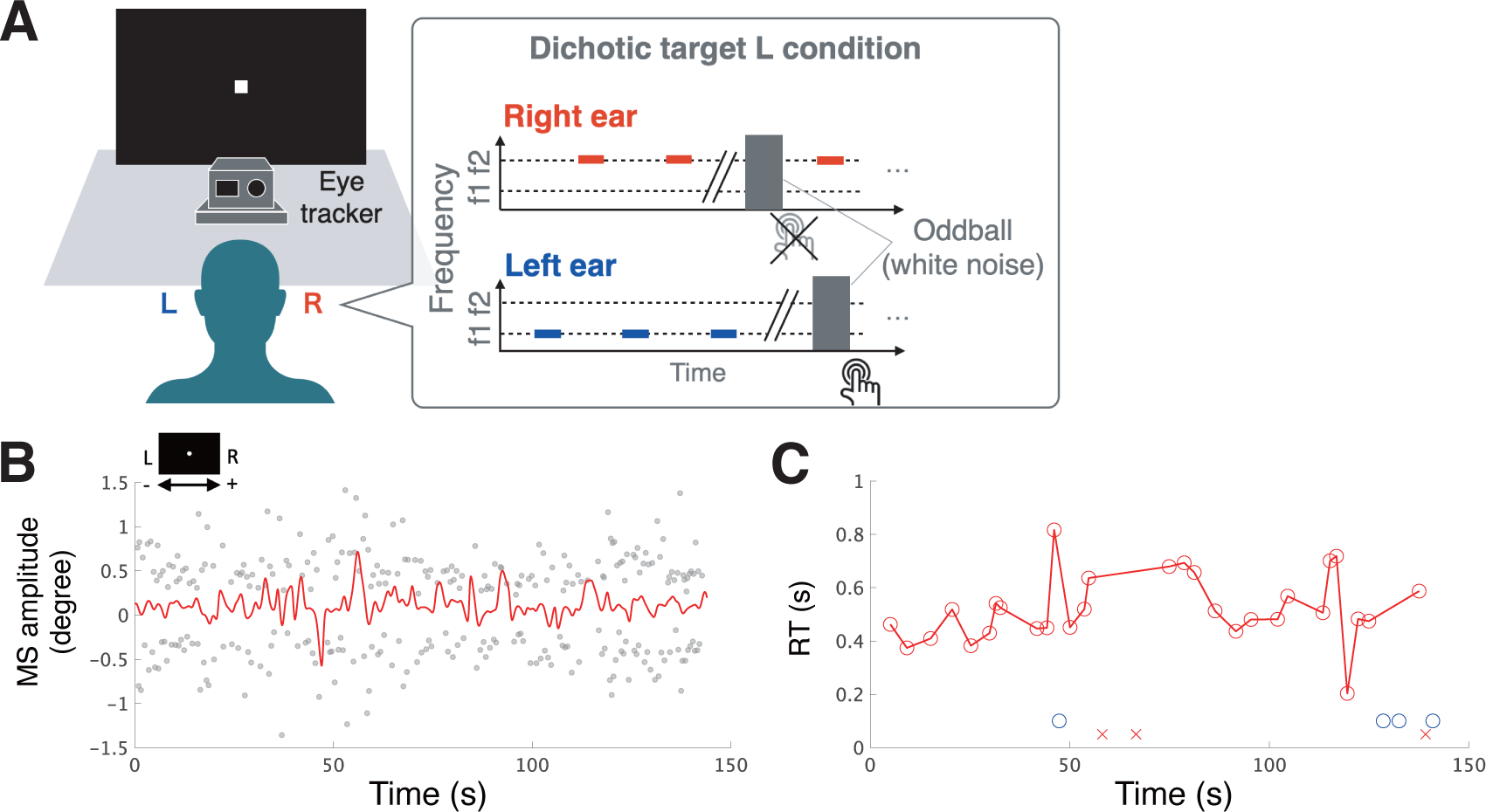
Experimental design. ***A***, Schematic illustration of a dichotic oddball detection task. The participants were asked to detect oddball sounds (white noise) embedded in a sequence of standard sounds (complex tones) with alternating fundamental frequency (F0), presented to the left and right ears. This figure illustrates the Dich_TargetL condition, which is only shown as an example. Please see Table 1 for the summary of experimental conditions. We measured the behavioral responses by button press and the eye metrical information (gaze position and pupil size) using a camera-based eye tracker (Eyelink). In Experiments 1 and 3, we measured EEG to evaluate auditory brainstem frequency-following responses in addition to the eye metrics (not shown). ***B***, Example of the time course of microsaccade (MS) amplitudes for one participant’s data from Experiment 2 (only showing the Dich_TargetL condition with medium oddball intensity). The grey dots represent individual MS amplitudes. The red line represents the running-averaged value of the amplitudes. MSs are discrete events that occur randomly, and their amplitude fluctuates over time. ***C***, Example of the time course of reaction time (RT) of the oddball detection (the data correspond to the data shown in panel ***B***). Red circles represent the RTs obtained for responses to random-timed oddballs in one session. Red crosses indicate the trials where the participant failed to respond to the target (miss). Blue circles indicate the trials where the participant responded to the oddball sounds presented to the unattended ear (false alarm). The number of oddball sounds was 64 (32 for each ear) in one session for Experiment 2. RT also fluctuates over time, sometimes reaching around 1 s, indicating that the participant’s attention state varied during the task.

**Table 1.**
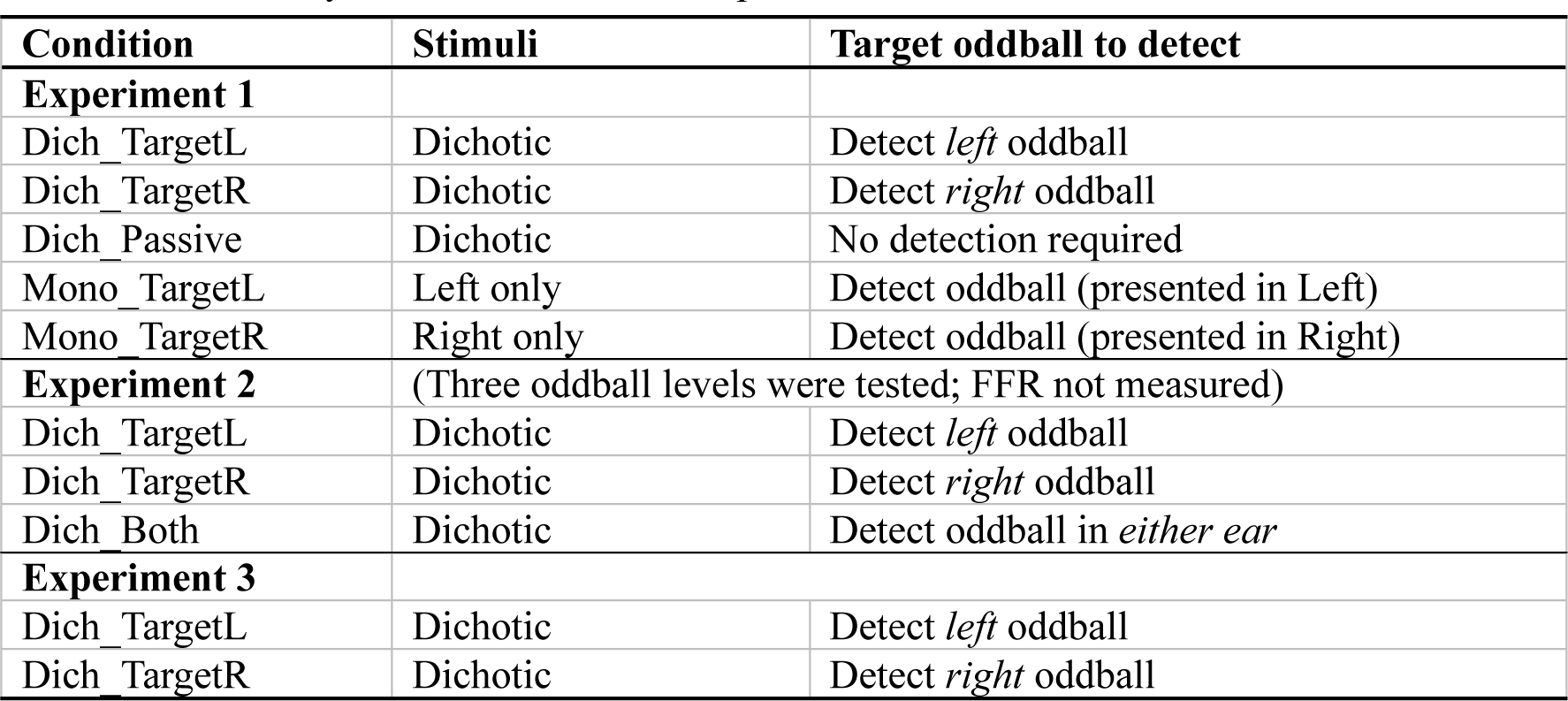
Summary of tasks for the three experiments.

| Condition | Stimuli | Target oddball to detect |
| --- | --- | --- |
| **Experiment 1** |  |  |
| Dich_TargetL | Dichotic | Detect *left* oddball |
| Dich_TargetR | Dichotic | Detect *right* oddball |
| Dich_Passive | Dichotic | No detection required |
| Mono_TargetL | Left only | Detect oddball (presented in Left) |
| Mono_TargetR | Right only | Detect oddball (presented in Right) |
| **Experiment 2** | (Three oddball levels were tested; FFR not measured) |  |
| Dich_TargetL | Dichotic | Detect *left* oddball |
| Dich_TargetR | Dichotic | Detect *right* oddball |
| Dich_Both | Dichotic | Detect oddball in *either ear* |
| **Experiment 3** |  |  |
| Dich_TargetL | Dichotic | Detect *left* oddball |
| Dich_TargetR | Dichotic | Detect *right* oddball |

Experiment 1 included five conditions that differed in stimulus presentation and task: In the *Dich_TargetL* and *Dich_TargetR* conditions, the stimuli were presented dichotically, and the participant was asked to maintain attention toward the left or right ear only, respectively, and respond as quickly as possible only to the oddball sounds presented to that ear. In the *Dich_Passive* condition, although the stimuli were again dichotically presented, participants were asked to listen to the stimuli passively, i.e., not attending to a particular ear and not responding to oddballs. In the *Mono_TargetL* and *Mono_TargetR* conditions, the stimuli were presented to the left or right ear only, respectively, and the participant was asked to detect oddballs in that ear (thus no selection between the ears was required). In one block of five sessions (corresponding to the five conditions), the conditions were ordered pseudo-randomly. We conducted 18 blocks per participant in total.

In Experiment 2, the Dich_TargetL, Dich_TargetR, and *Dich_TargetBoth* conditions were investigated. The former two conditions were the same as in Experiment 1, while we tested three oddball levels (*soft*, *mid*, and *loud*) to examine the effect of the intensity of the oddball sound on MS. We considered that the oddball intensity would be associated with the salience of the oddball, as well as the difficulty of the task. In the Dich_TargetBoth condition, participants were asked to attend to both ears and respond to oddballs presented in either ear. This means that they were not required to facilitate or suppress responses to a particular ear. In one block of nine sessions (corresponding to the nine conditions, i.e., three attention directions times, three intensities), the conditions were ordered pseudo-randomly. We divided one block into three sub-blocks and ordered the conditions so that each sub-block contained the Dich_TargetL, Dich_TargetR, and Dich_TargetBoth conditions in a pseudo-random order. We conducted 6 blocks per participant in total.

In Experiment 3, we focused only on the Dich_Target L and Dich_Target R conditions to yield a large number of repetitions for the same condition within a limited time. This experiment was conducted to ensure the reliability of the scalp-recorded subcortical responses, which are susceptible to the number of trials.

In Experiments 1 and 2, standard sounds were harmonic complex tones with F0s of 315 Hz and 395 Hz, presented to the left and right ears, respectively. In Experiment 3, the F0s of the complex tones were 375 Hz and 470 Hz, and the presented ears were counterbalanced between participants. The number of harmonic components was five (including F0) for all experiments. The starting phase of the harmonic components was fixed across presentations. We chose these harmonic complex tones because FFR (regarded as originating from the brainstem) can be reliably obtained for F0s up to about 500 Hz (Hoormann et al., 1992; Tichko and Skoe, 2017a) while avoiding confounds caused by a slow cortical following response (Coffey et al., 2016; Tichko and Skoe, 2017a). The A-weighted sound pressure level of standard sounds was 78 dB, calibrated with an artificial ear with a coupler by IEC 60318–1. The oddball sound was white noise and presented in place of the standard. The duration of each standard and oddball sound was 40 ms and the interval between the sounds was 50 ms for Experiments 1 and 2. In Experiment 3, the duration of each sound was 50 ms and the interval between the sounds was jittered from 50 to 63 ms with uniformly distributed random numbers. In Experiment 1, the A-weighted sound pressure level of the oddball was the same as the standard sounds. In Experiment 2, we tested three intensities to vary the detectability of oddballs; 2.5, 5, and 10 dB relative to the detection threshold of the oddballs measured prior to the main experiment for each participant. In Experiment 3, the intensity of oddball sounds was the same as the 5-dB condition in Experiment 2. Thus, the saliency of oddball sounds was relatively low compared with Experiment 1. The total number of sounds, including the standards and oddballs, in Experiments 1 to 3 was 500, 800, and 700, respectively. Of these, the oddballs made up 4%, 8%, and 2%, respectively.

For all experiments, we presented a visual fixation spot on the center of the display (0.63 cd/m^2^). We wanted to investigate the relationship between FFR and baseline pupil size, which may reflect the participant’s arousal state, as well as microsaccades. The pupil size, however, reflects not only arousal states but also the brightness of visual stimuli. Thus, background luminance was manipulated only for Experiment 1 to dissociate the effects of brightness on baseline pupil size: For half sessions, 10.15 cd/m^2^, and for the other half, 2.80 cd/m^2^. For Experiments 2 and 3, the background luminance was 10.15 cd/m^2^ throughout the experiment.

### Detection of microsaccades

We analyzed the gaze data as the eye-tracker output for both eyes. To detect microsaccades (MSs), we defined the onset as the time when the velocity of gaze movement exceeded a certain threshold. Specifically, we used a threshold that was six times the standard deviation of the velocity in one trial (Engbert and Kliegl, 2003) We excluded large saccades whose amplitude exceeded 1.5 degrees in visual angle to extract only microsaccades. In the present study, we only focus on binocular MSs (Engbert and Kliegl, 2003), defined by a temporal overlap between the left-eye and right-eye MSs: if the difference between the onsets of left and right MSs exceeded 10 ms, we excluded those events from the analysis. We extracted only the horizontal component of MSs because we focused on the attention effect, which should be observed in the horizontal direction (i.e., left or right). Raw gaze positions were converted from the pixel values obtained by a five-point grid calibration into degrees of visual angle using the distance from the center of the display to the participant’s eyes. We excluded the eye data during blinks and 100 ms before the onset and after the offset of blinks and unsuccessful recordings from the analysis. These criteria for excluding data are common to all analyses related to eye metrical measures (i.e., gaze or pupil analysis).

### Analysis of microsaccades

We calculated the ratio of rightward microsaccade counts throughout a trial (Right MS ratio: # *MS*_*s*_⁄#*MS*_*all*_) to assess the overall bias of MS during an attention task. We also analyzed the peri-target amplitude time course of MS to examine the transient impact of the oddball (i.e., target) presentation on MS. To calculate the time course, the amplitudes of MSs that occurred within the -2 s to 2 s time range around each target presentation were found. Over multiple presentations of targets for a condition of interest, the amplitudes at discrete occurrence times relative to the target were averaged with a moving Gaussian window (*σ* = 100 ms, duration = 800 ms). We applied a cluster-based permutation test which has been commonly used for EEG and magnetoencephalographic (MEG) data (Maris & Oostenveld, 2007) for comparison of time courses of MS amplitude around oddball presentations to evaluate if there were specific time points when MSs occurred toward or away from the stimulus.

### Analysis of behavior

We evaluated moment-by-moment behavioral performance for target detection in terms of reaction time and correct-rejection or false-alarm response. We sought button response events with a time window from 0.15 s to 1s after oddball presentations. For example, if the participant failed to press the button within 1.5 s after an oddball, that trial was excluded from the analysis as a “miss” trial. False alarm trials were defined as when participants incorrectly responded to the oddball sound presented to an unattended ear. We used a relatively long time window for the false alarm detection (0.15 s to 2 s from the oddball) to capture deterioration in the level of alertness and to ensure we had sufficient trial numbers. The analyses of false alarm trials were conducted only for the Dich_TargetL condition or Dich_TargetR condition.

### Analyses of subcortical activity (FFR)

We first applied the bandpass filter (passband: 70 – 2000 Hz) to the raw EEG signal. EEG recordings corresponding to each presentation of a standard sound were extracted using a rectangular window. If the root-mean-squared amplitude of the segmented signal exceeded 35 µV, that extract was excluded from further analyses (Skoe and Kraus, 2010). Then, the amplitude spectrum of the extracted segment was calculated by fast Fourier transformation (FFT) with a 40-ms Hanning window. Finally, we derived FFRs as the amplitude of the spectral component of the EEG signal at the F0 of the standard sound by averaging the FFR value for all extracted segments (Krizman and Kraus, 2019).

### Peri-MS FFR functions

We also examined the relationship between MS and FFR in an MS-by-MS manner. The general procedure is illustrated in Fig. 9A. First, for a given MS towards the direction of interest, standard sounds (separately for the left- and right-ear sounds) within 0.5 s before and after MS onset were found, and EEG responses to the sounds were fast-Fourier-transformed. We extracted the spectral components as complex numbers only at the stimulus F0 frequency. Both the real and imaginary parts (i.e., amplitude and phase information) were necessary to improve sensitivity to the stimulus-related response by focusing on the phase-locked component relative to the stimulus waveform. The extracted values were represented as point data, expressing the FFR strength (in complex numbers) at times relative to the MS. We computed the FFR amplitude not only during the stimulus presentation but also during the pre-stimulus period (i.e., the silent period between -40 and 0 ms relative to the sound onset) to evaluate the baseline level of FFR for each participant. If the adjacent MS occurred within 500 ms before or after the target MS, that trial was excluded from the analysis because the adjacent MS may affect the FFR time course. Next, a moving Gaussian window (s = 50 ms, duration = 400 ms) along the time axis was used to average the pooled data across the MS occurrences (Fig. 9A). After averaging, data were transformed into absolute values (spectral amplitudes) and grand averaged across participants. The obtained smooth functions are referred to as peri-MS FFR functions, which is the time course of FFR amplitude around the MS.

We derived (normalized) peri-MS FFR functions for the two cases when the MS direction was congruent and incongruent with the stimulus ear for which FFR was evaluated (referred to as the congruent- and incongruent-MS cases, respectively), summarizing the data across all the participants (after excluding a few participants with poor signal-to-noise ratio; see below) and for the Dich_TargetL and Dich_TargetR conditions. The steps of this process are as follows. First, we calculated the difference between the peri-MS FFR function derived from the stimulus presentation time and that derived from the pre-stimulus time (baseline-corrected peri-MS FFR function). Second, the amplitude of the baseline corrected function was expressed as a z-score to disregard the effects of differences in FFR amplitude (nV) that depended on participant and stimulus frequency. It should be noted that in the above process, we excluded the data from two participants (Experiment 1) and one participant (Experiment 2), whose signal-to-noise ratio (SNR) was smaller than 0 dB (FFR to the stimulus was on average smaller than the baseline level during no stimulation). The SNR was defined as the ratio of the averaged spectral amplitude of FFR at the stimulus frequencies during the stimulus presentation to that during the pre-stimulus time. Poor SNRs for the excluded participants can be explained partly by the relatively small number of MSs, which determined the number of samples to be averaged.

## Results

### Stimulus-related microsaccade bias: MS amplitude systematically varied after the oddball presentation

We first asked whether the MS amplitude shows systematic change after the presentation of oddball sounds depending on the stimulus direction, as shown in the previous visual studies (see Hafed et al., 2021 for review). To describe the MS bias referenced to the oddball direction, we defined “oddball-referenced MS bias” by subtracting the trace of MS amplitude around the left-oddball presentation in the Dich_TargetL condition (the left columns of Fig. 2 A and B, blue lines) from that around the right-oddball presentation in the Dich_TargetR condition (red lines), as shown schematically in the upper part of right panels in Figs. 2A and 2B. The *positive* value in this measure indicates the MS bias *toward* the direction of oddball sounds. MS amplitude was generally biased away from the oddball sound around 1 to 2 s after the oddball presentations in the Dich_TargetL/R condition, which was commonly observed in both Experiments 1 and 2 (right columns in Figs. 2A and 2B). Interestingly, for the Dich_TargetBoth condition in Experiment 2, we found an early MS bias toward the oddball followed by a bias away (lower right panel in Fig. 2B). The horizontal grey lines in the left panels indicate clusters with significant differences between left- and right-oddball presentations. The horizontal red and blue lines represent clusters with significantly different values from the baseline, while the lines above and below represent clusters with greater or smaller values compared to the baseline, respectively. The baseline was derived by averaging MS amplitude in the time range of -0.5 to 0 s from the oddball onset (represented by dotted lines with corresponding colors). The black horizontal lines above and below in the right panels indicate clusters with greater or smaller values compared to zero for the oddball-reference MS bias, respectively.

**Figure 2.**
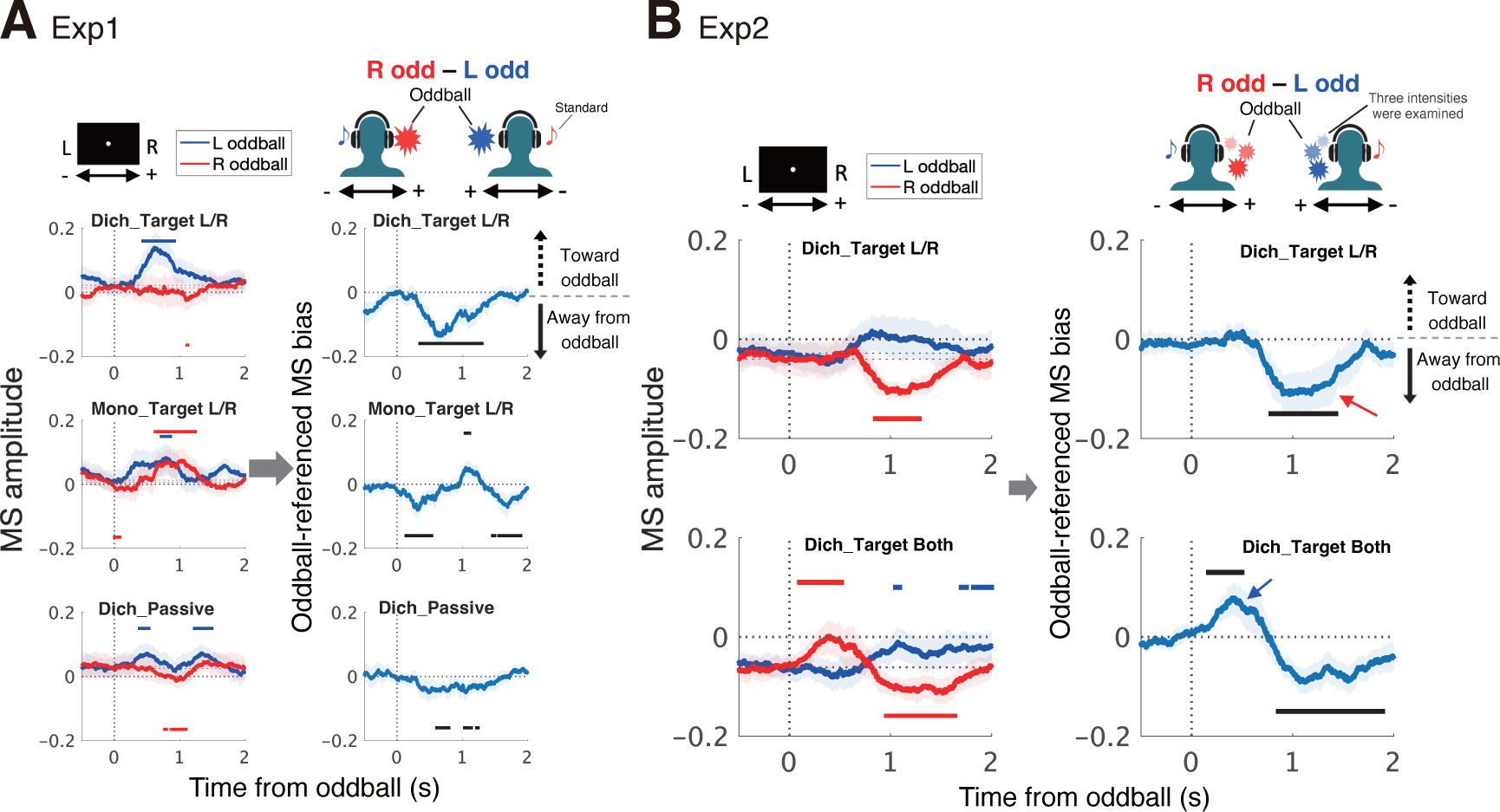
MS bias around the oddball presentation. ***A***, MS bias for Experiment 1. The detected MSs were averaged using a moving Gaussian window (see, Methods). We focused on the MS amplitude only during the oddball presentation, from -2 to 2 s relative to the oddball onset. As shown in the schematic illustration at the top left, positive and negative values of MS amplitudes in the left panels indicate rightward and leftward eye movements, respectively. The schematic illustration in the upper right depicts the relationship of sign between the oddball location and the MS direction. Blue and red horizontal bars represent the significant clusters that differed from the baseline amplitude, derived from the range -0.2 to 0 s from the oddball onset, for the Dich_Target L and R conditions, respectively. Horizontal bars above and below represent the clusters exhibited rightward and leftward MS shifts compared to the baseline, respectively. The right panels show the oddball-referenced MS bias (see, Results). Black horizontal bars represent the significant clusters that differed from zero, where positive and negative values indicate the direction toward and opposite to oddballs, respectively. We found a tendency whereby MS biased away from the oddball sounds in the Dich_Target L/R conditions. ***B***, MS bias for Experiment 2, where the intensity of oddball sound was controlled to make the task difficult compared with Experiment 1. Essentially the same conventions as A (Experiment 1) are used to represent the data. The figures show averaged results across multiple sound levels. The result shows that the MS was biased away from the oddball direction around 1 s after the oddball presentation (see the red arrows in the right panel), as observed in Experiment 1. We found the early MS bias (∼0.5 s) toward the oddball only for the Dich_Both condition (the blue arrow), where the participants were not required to perform selective detection between left-ear and right-ear sounds.

### Stimulus-related microsaccade bias associated with task performance in a dichotic selective detection task

We next asked to what extent the MS reflects the task performance, as an index of attention states that fluctuate over time. The analysis was inspired by the fact that the MS toward the oddball sounds was observed for the Dich_TargetBoth condition but not for the Dich_TargetL/R conditions. The Dich_TargetL/R conditions require a selective response to left or right ears, respectively, while a task in the Dich_TargetBoth condition requires simple reactions, i.e., no need to suppress responses to sounds in the non-target ear. We hypothesized that such a selective reaction process would somehow be reflected in MS bias: The early reflexive MS toward the stimulus (orienting MS, see Hafed et al., 2021) would be suppressed during the dichotic selective detection task to reduce the risk of reflexively responding to the oddballs. More specifically, we expected that the orienting MS would occur toward the oddballs when the participant failed to inhibit the orienting ocular response toward the salient sound even during the Dich_TargetL/R conditions, due to, for example, the participants’ low alertness level at that moment.

Under this hypothesis, we first compared the oddball-referenced MS bias between correct-rejection (CR) and false-alarm (FA) trials. The CR and FA trials were defined as the trials where participants correctly suppress the response or incorrectly respond, respectively, to the oddballs presented to the unattended ear. The false alarm rate was significantly greater for louder oddball sounds in Experiment 2 (right panel in Fig. 3A), assuring that the false alarm responses were associated with the salience (i.e., difficulty in ignoring) of the sound in the unattended ear. We observed that the oddball-referenced MS bias exhibited positive values during FA trials around oddball presentations for both Experiments 1 and 2 (Fig. 3B). This may indicate that the MS reflects endogenous factors that influence the performance of the selective listening task. Interestingly, such FA-related MS bias occurred in concurrence with or even before the timing of the oddball presentation, indicating that the MS bias was not triggered by the stimuli but was related to the internal state: The MS coincidentally occurring toward an unattended oddball may predict an FA response.

**Figure 3.**
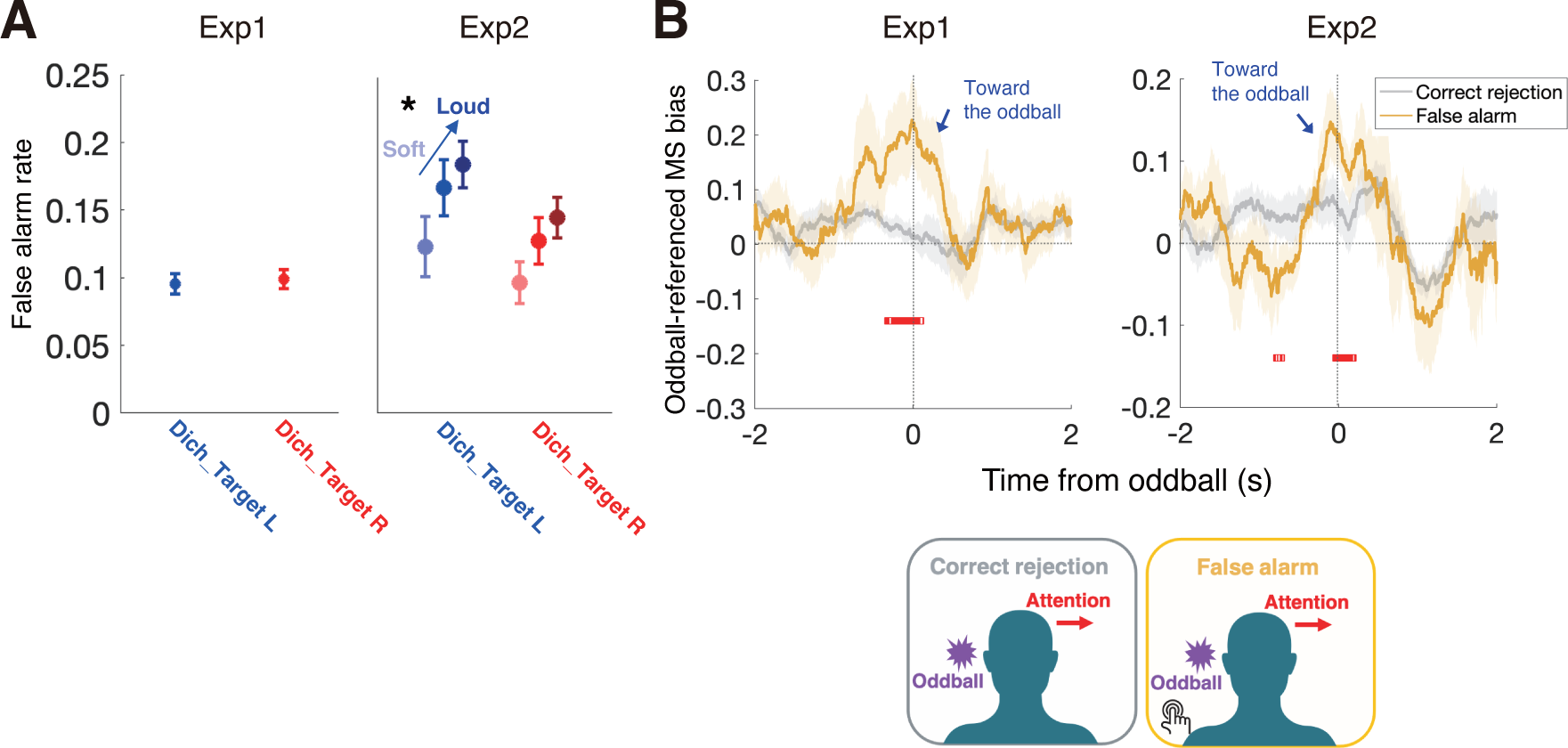
MS bias toward the oddball occurred for false alarm trials. ***A***, False alarm (FA) rates for Experiments 1 and 2. The deviation of FA rates was generally larger for Experiment 2 than for Experiment 1, probably reflecting the task difficulty. FA rate was highest for the highest intensity condition in Experiment 2. ***B***, MS bias around oddballs for FA trials and for correct rejection (CR) trials (yellow and gray lines, respectively). We found significant clusters (shown by red squares) for both Experiments 1 and 2 around the timing of the oddball presentation.

Second, we focused on reaction time (RT) as another indicator of alertness level or performance. Generally, the RT was significantly shorter for the Mono_TargetL/R conditions and the Dich_TargetBoth condition (simple reaction tasks) than for the Dich_TargetL/R conditions in Experiments 1 and 2, respectively (Fig. 4A). These results indicate that the Dich_TargetL/R conditions required additional processing load for selective detection, compared with the Mono_TargetL/R and Dich_TargetBoth conditions.

**Figure 4.**
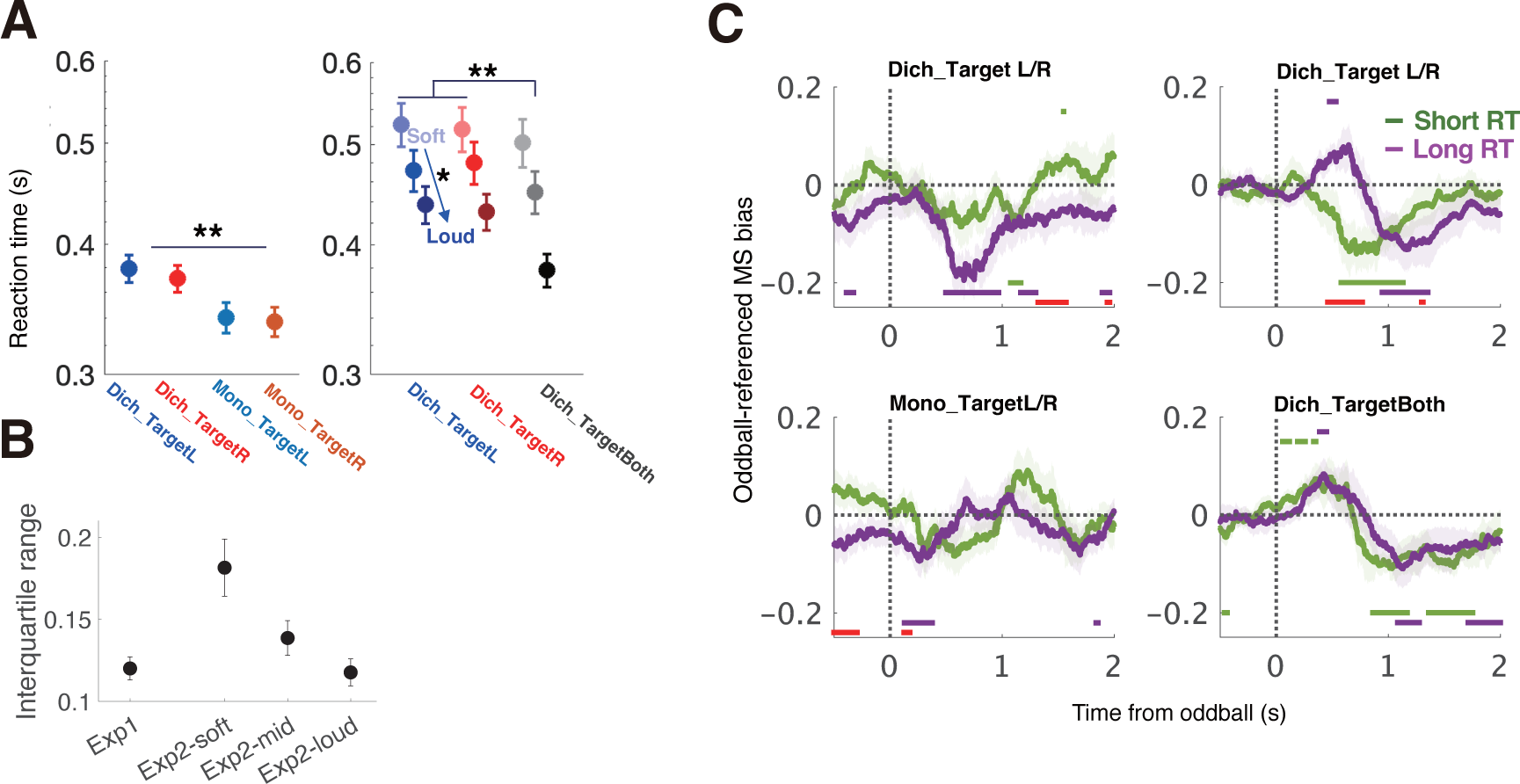
Orienting MS occurred for long RT trials but not for short RT trials. ***A***, Mean reaction time (RT) for Experiments 1 and 2. RT was significantly longer for the Dich_TargetL/R conditions than for the monoaural condition in Experiment 1 or the Dich_TargetBoth condition in Experiment 2. We found a significant effect of oddball intensity on RTs (long and short RTs for soft and loud oddballs, respectively). ***B***, Variation in RTs (interquartile range) for Experiments 1 and 2. ***C***, Oddball-referenced MS bias for the short and long RT trials. We found early MS bias *toward* the oddball sounds in long-RT trials for the Dich_TargetL/R conditions in Experiment 2 (i.e., low performance), not observed in short-RT trials (i.e., high performance). The temporal pattern of the MS bias function was similar to that for the Dich_TargetBoth condition, suggesting that the orienting MS tends to occur when participants’ alertness level is low. In Experiment 1, there was no clear evidence of the association between RT and MS bias, probably because RT variation was smaller than in Experiment 2 (panel ***C***) and was insufficient to observe its effect on MS.

Does MS bias reflect trial-by-trial variation in RT during the Dich_TargetL/R task? To answer this question, we divided the trials of each condition and participants into halves with RTs shorter and longer than the median (referred to as short- and long-RT trials, respectively). We found that early MS bias *toward* the oddball sounds in long-RT trials for the Dich_TargetL/R conditions in Experiment 2 (i.e., low performance) was not observed in short-RT trials (the right upper panel in Fig.4C). We found no significant effect of RT on oddball-directed MS bias for the Dich_TargetL/R conditions in Experiment 1, implying that the task in Experiment 1 was relatively easy compared with Experiment 2 and thus the RTs were generally short (Fig. 4A) and had a small variance (Fig. 4B). The small variation in RT in Experiment 1 probably made it difficult to reveal its correlation with MS bias. We also found no significant effect of RT on MS bias for the Dich_TargetBoth condition in Experiment 2. This may reflect the fact that, unlike the Dich_TargetL/R conditions, the Dich_TargetBoth condition did not require selective attention that needs inhibition of the orienting responses toward the salient sounds, which could be a distractor in the dichotic sound detection task.

In summary, we consistently found that the early MS bias toward the oddballs occurred when the task performance was low (i.e., false-alarm or long-RT trials) or for the Dich_TargetBoth condition. These results lead us to infer that the orienting MS toward the salient stimulus tends to occur when the participants’ alertness level is low during a dichotic selective attention task and when they engage in a simple reaction task to respond to targets in the dichotic sound sequence, which requires no selective attention between ears.

### Microsaccade reflects the direction of sustained auditory attention

We next asked whether the MS direction was biased toward or away from the direction of voluntary and sustained auditory attention, regardless of the oddball presentation (see Xue et al., 2020 for the visual domain). The MS bias was quantified by computing the *rightward MS ratio*, the number of rightward MS divided by the total number of MS for each condition and participant. There was a tendency that the rightward MS ratio would be higher when the participant attended to the left ear (Dich_TargetL) than when they attended to the right ear (Dich_TargetR) in both Experiments 1 and 2 (Fig. 5A; comparison between the blue and red symbols). This relationship between the directions of attention and MS was opposite to the one observed for the visual sustained attention task (Xue et al., 2020).

**Figure 5.**
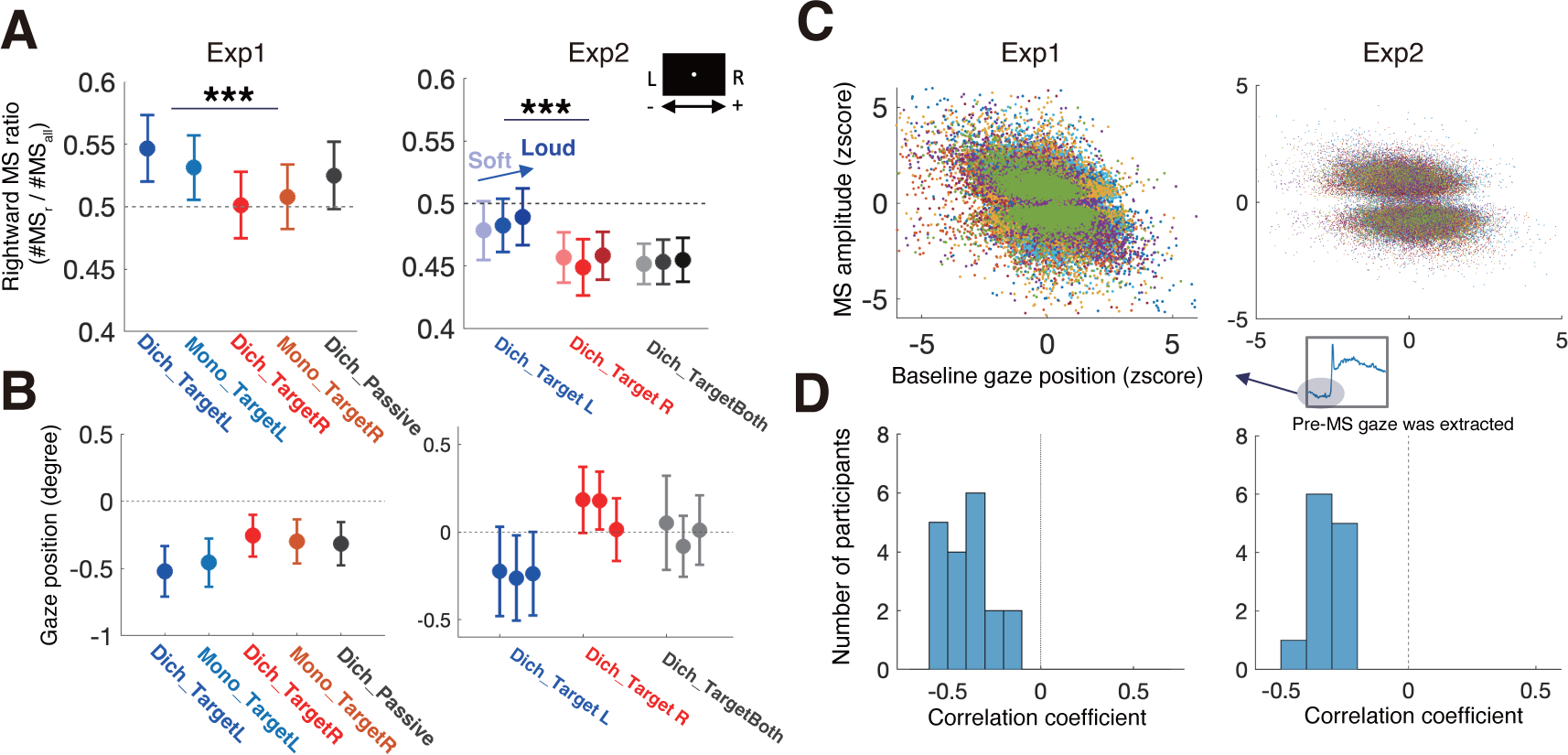
Effect of sustained attention and gaze position on MS bias. ***A***, Mean rightward MS ratio (the number of rightward MS was divided by the total number of all MSs). For both Experiments 1 and 2, MS tended to be biased away from the target side (e.g., the ratio of rightward MS was greater for the Dich_TargetL condition than for the Dich_TargetR condition). ***B***, Mean gaze position. A positive value of gaze position indicates a rightward gaze. The overall patterns of the gaze positions were mirror-symmetric to the patterns of the rightward MS ratio. ***C***, The relationship between the baseline gaze position and MS in z-score. Each point corresponds to one MS occurrence. The color of the dots represents the participant. ***D***, The distribution of correlation coefficients for the gaze position and MS amplitude, derived from individual participants. All participants showed a negative correlation, implying that the MS away from the target included the gaze-correcting MS.

As a statistical analysis of Experiment 1, we conducted two-factor repeated-measures ANOVA for rightward MS ratio in the Dich_TargetL, Dich_TargetR, Mono_TargetL, and Mono_TargetR conditions. The factors were the presentation method (monaural versus dichotic) and target laterality (left-versus right-ear targets). We found a significant effect of laterality (p < 0.00001, F = 40.509, left panel in Fig. 5A, blue > red symbols) showing the MS bias away from the target direction. We found that the presentation method had no significant effect (p = 0.494). This indicates that the MS bias away from the target direction occurs even when selective attention is not required (Mono_TargetL/R conditions).

It is interesting that the rightward MS ratio, when pooled across all the conditions, was generally distributed above 0.5, indicating that the MS direction was biased towards the right, regardless of the target side. Note, however, that the ratio for the Dich_Passive alone, where the listener was not required to pay attention to the stimuli, was not significantly above 0.5 (p = 0.3680).

We conducted the same analysis for Experiment 2; ANOVA for rightward MS ratio in the Dich_TargetL and Dich_TargetR conditions. In this case, the factors were the target laterality (left and right) and sound intensity (soft, middle, and loud). We found that the rightward MS ratio was larger for the Dich_TargetL condition than for the Dich_TargetR condition (p = 0.0020, F = 16.303, Fig. 5A right panel, blue > red symbols), indicating that the MS was biased away from the target direction, a finding consistent with Experiment 1. We found no significant effect of the sound intensity (p = 0.293) and no significant interaction between the two factors (p = 0.619). This time, the rightward MS ratio was generally below 0.5, when pooled across the three conditions (i.e., Dich_TargetL, Dich_TargetR, and Dich_TargetBoth) or when focused on only the Dich_TargetBoth condition (black symbols in the right panel of Fig. 5A); p=0.0867 or 0.0183, respectively, by one-sample t-test. We have no explanation for this discrepancy in the general bias between Experiments 1 and 2.

### MS bias reflecting sustained attention can be explained by the spontaneous gaze shift

MS bias away from the target side may be accounted for, at least in part, by corrective saccades for an error between the fixation dot and the gaze position (Costela et al., 2014), which can be biased toward the direction of sustained attention (Gopher, 1973). To confirm this, we sampled the gaze position data at the time immediately before the MS onsets. Therefore, all gaze data in this analysis were linked with the MS timing. As expected from Gopher (1973), the averaged gaze direction was biased toward the attended direction (Fig. 5B). The overall patterns of the gaze positions (Fig. 5B) appeared to be mirror-symmetric to the patterns of the rightward MS ratios (Fig. 5A). That is, in conditions with higher rightward MS ratios, for example, the gaze tended to be towards the left side. The trial-by-trial correlation between the gaze amplitude and the MS amplitude including direction information (rightward; positive, leftward; negative) for each participant separately (Fig. 5C) exhibited a consistent tendency, i.e., negative coefficients for all participants (Fig. 6D). This indicates the compensatory relationship between MS and gaze amplitudes and is consistent with the above explanation by corrective saccades. This result suggests that the MS away from the target direction included the gaze-correcting MS due to the spontaneous gaze shift toward the direction of sustained auditory attention.

**Figure 6.**
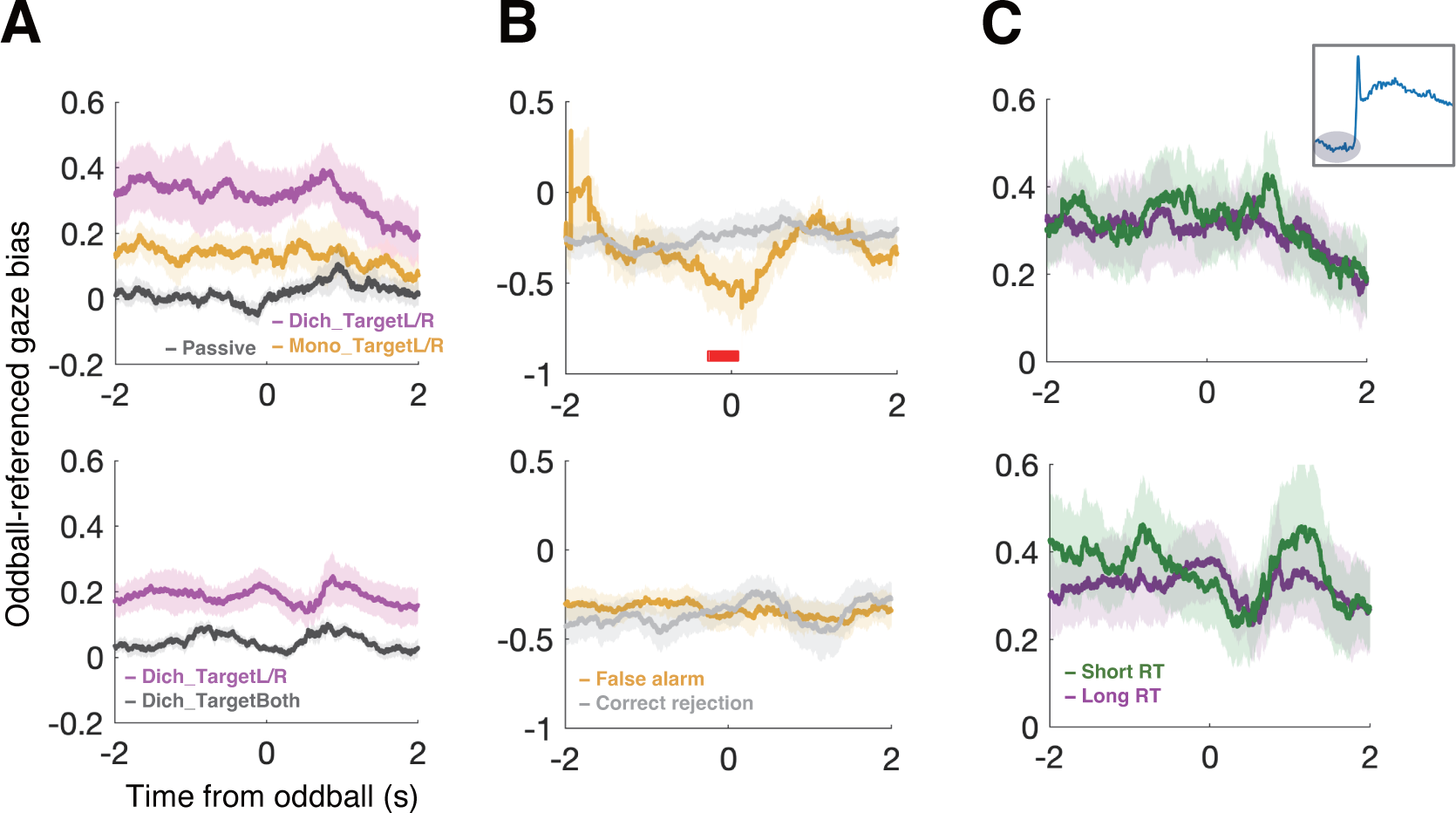
The gaze time course around the oddball presentation did not show a dependence on task performance. Gaze positions were extracted from the time immediately before the MS onset. Thus, these data did not include gaze changes due to MS. ***A***, Oddball-referenced gaze bias. The gaze position was biased toward the target side for the Dich_TargetL/R conditions (magenta line) or the Mono_TargetL/R conditions (yellow line). ***B***, The time course of gaze bias around the oddball sound for FA and CR trials. ***C***, The time course of gaze bias around the oddball sound for short-RT and long-RT trials. No clear relationship between gaze bias and task performance was observed except for the difference between FA and CR trials in Experiment 1.

### The gaze shift, however, cannot account for the link between the stimulus-related MS bias and task performance

One may doubt that the stimulus-related MS bias described in earlier sections (Fig. 2-4) might be influenced by the gaze modulation due to the oddball presentation which, in turn, possibly affects MS direction in a compensatory manner as shown in Fig.6. We addressed this concern by calculating the oddball-referenced *gaze* bias in the same way as the analysis of oddball-referenced MS bias (Fig.2). Unlike the MS, there were no specific patterns in gaze bias around oddball presentations (Fig. 6A). The overall positive bias of the oddball-referenced gaze is a replication of the observations in Fig. 5B that the gaze direction was shifted toward the direction of sustained attention. Comparing low- and high-performance trials (i.e., FA and CR trials, short- and long-RT trials), we found no significant difference between FA and CR trials in the gaze time course for Experiment 2 and no significant effect of RT on the gaze time course (Fig. 6C). These results indicate that although the MS bias generally correlated with the gaze position in a compensatory manner (Fig. 5), the oddball-induced effect on MS direction associated with the task performance of an auditory selective attention task cannot be explained by gaze drift. However, there was a period with a significant difference in the gaze position between the FA and CR trials for Experiment 1 (Fig. 6B, upper panel), which was close to that of the MS result (Fig. 3B). This momentary gaze shift might have contributed to the oddball-reference MS bias, although the reason for the discrepancy between Experiments 1 and 2 is unclear.

### Microsaccade amplitude correlates with baseline pupil size on a large time scale, but not on a trial-by-trial basis

Can the observed relationship between task performance and MS bias be accounted for by the arousal state? We addressed this question by examining the baseline pupil size during an attention task. The baseline pupil size is known to reflect an arousal state linked to the locus-coeruleus norepinephrine system (LC-NE system, Aston-Jones & Cohen, 2005). We divided the MS data according to pupil size (small, mid, and large). We found a significant tendency that the target-referenced MS bias was decreased with increased pupil size for Experiment 1, but not for Experiment 2 (upper panels in Fig. 7C, one-factor repeated measures ANOVA; p = 0.0391 for Experiment 1 and p = 0.577 for Experiment 2). This tendency was largely due to the long-term trend of pupil size and MS amplitude within one experimental session (Fig. 7B), i.e., the first 100 trials exhibited the greatest MS bias and baseline pupil size, which may reflect the cognitive fatigue or neural adaptation correlated with the saccade and pupil systems independently. Then, we evaluated the trial-by-trial association between the two measures by removing this slow trend. The trend of changes in pupil size and MS amplitude was derived by smoothing with a large temporal window (100 trials) throughout one session (examples of the analysis of pupil size changes in one session are shown in Fig. 7A). We next examined the relationship between the trial-by-trial changes in pupil size and MS amplitude after subtracting the trend component. As a result, the correlation between the MS bias away from the target and pupil size was no longer observed (lower panels in Fig. 7C, p = 0.658 and p = 0.863 for experiments 1 and 2, respectively), suggesting that the relationship between pupil size and MS reflects global changes in neural activity or adaptation on a large time scale, not trial-by-trial fluctuations in arousal states.

**Figure 7.**
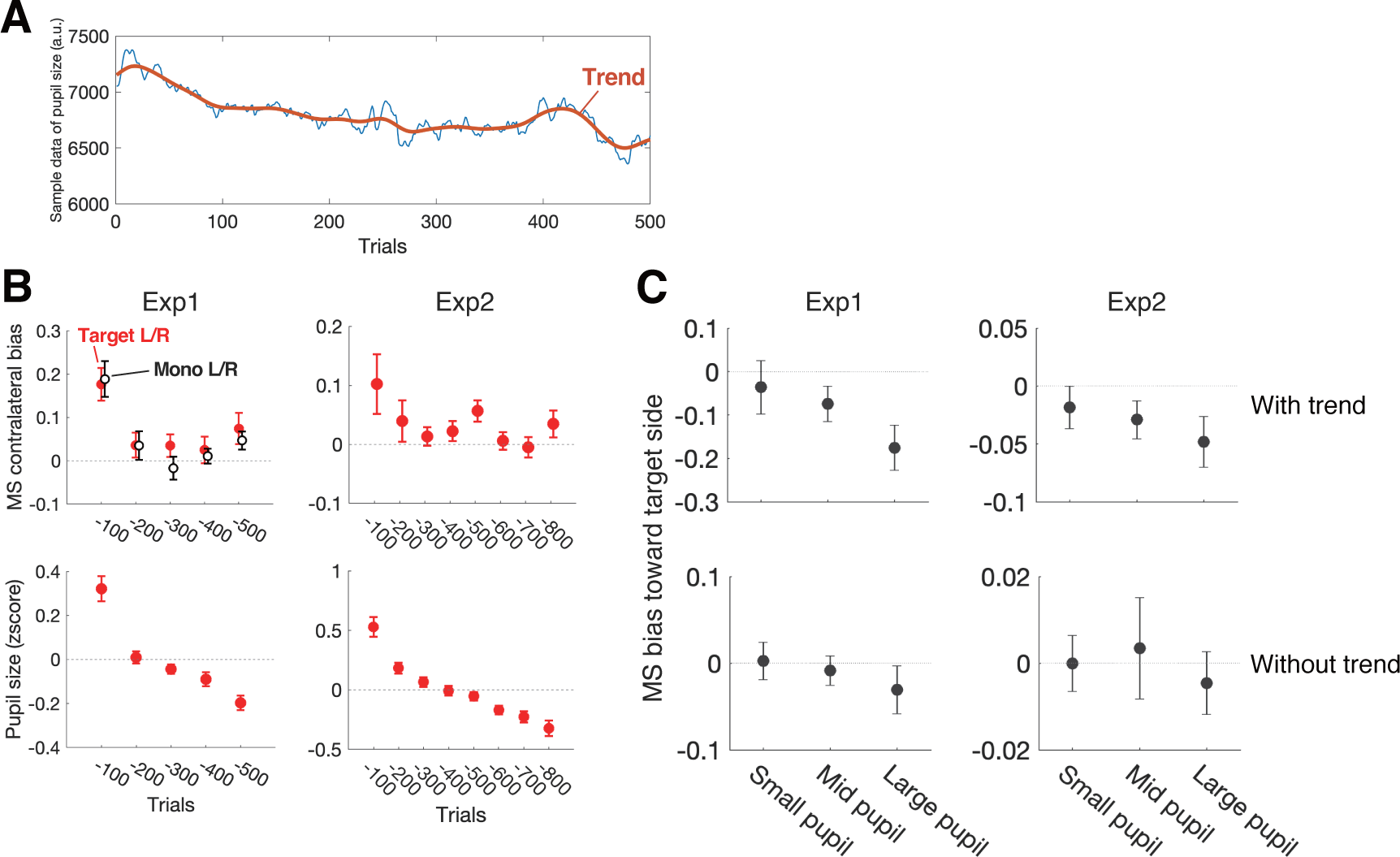
Relationship between MS and pupil size. ***A***, An example of the time course of pupil size from the data obtained by Experiment 1, showing that it gradually decreased over the long term with short-term fluctuation. ***B***, The trend of long-term changes in MS bias and pupil size for Experiments 1 (left panels) and 2 (right panels). Both MS bias away from the oddballs and pupil size exhibited large values for the first 100 trials. Pupil size changes showed a decreasing trend throughout the experimental session. ***C***, Relationship between MS and pupil size with and without trend components. MS bias away from the oddball sounds tended to be greater for large pupil size than for small pupil size (upper panels). If we excluded the trend component of pupil size change throughout the experimental session, the relationship between MS bias away from the oddballs and pupil size was no longer observed (lower panels).

### Auditory subcortical response (FFR): No significant effect of the attentional set

Finally, we asked whether the auditory processing (specifically, brainstem frequency-following response: FFR) co-varies with the attentional set and/or MS bias during the dichotic oddball detection task. Figure 8A shows bandpass-filtered EEG responses (i.e., FFRs) to a pair of consecutive left-ear (315 Hz) and right-ear (395 Hz) sounds, averaged across all conditions in Experiment 1. We measured the FFRs, which are responses corresponding to the stimuli in time (with some delay) and frequency. We can identify responses to the left- and right-ear sounds by referring to the EEG energy at the frequencies of the respective sounds. Thus, the FFR amplitudes for the left- and right-ear sounds were defined as the spectral EEG amplitudes at 315 and 395 Hz, respectively, and are plotted in Fig. 8B.

**Figure 8.**
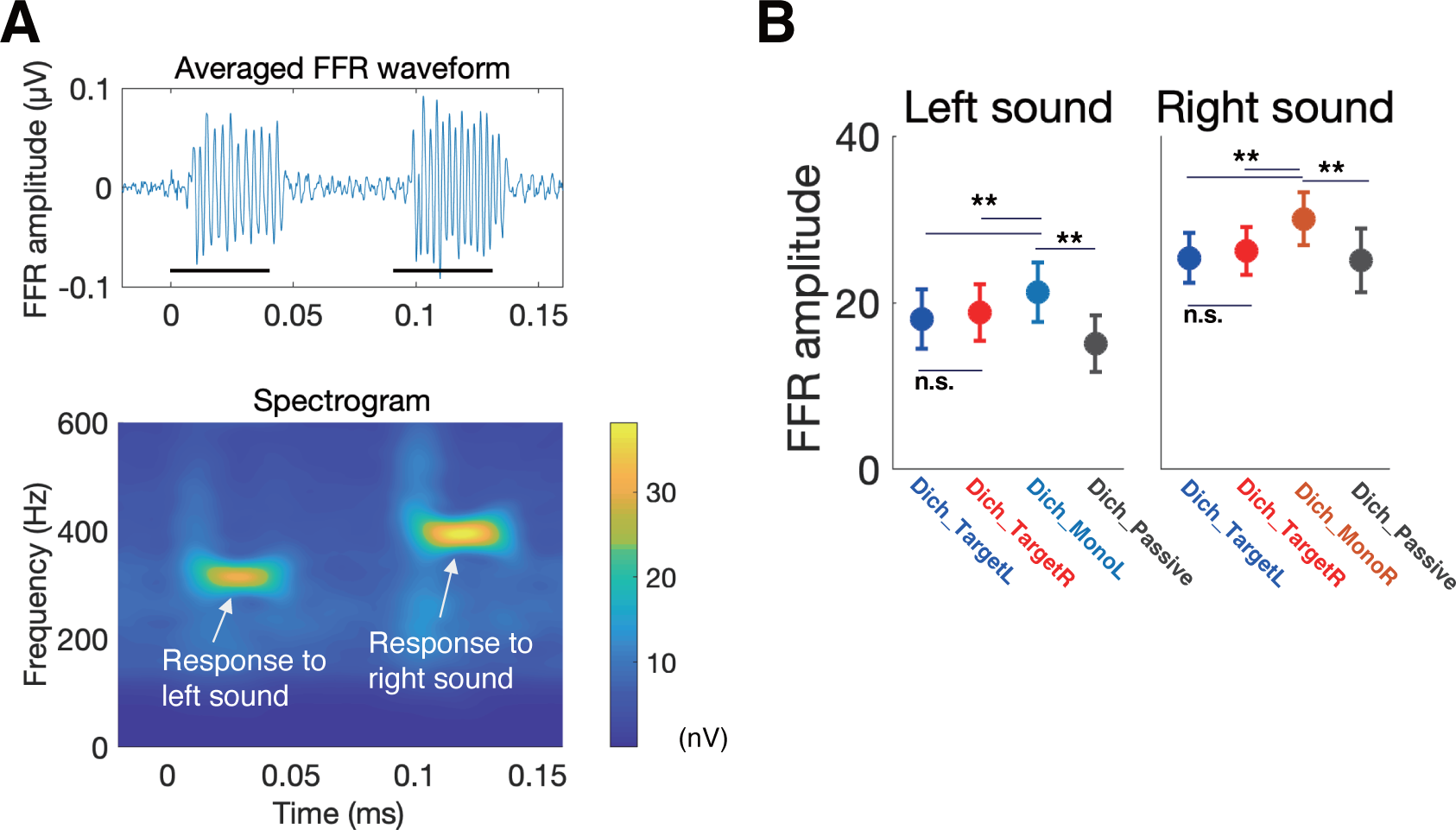
No significant effect of dichotic attention on average FFR strength. ***A***, Grand-averaged waveform and spectrogram of FFR obtained in Experiment 1. The black bars under the waveform indicate the duration of stimulus presentation. The spectrogram shows the power at the frequencies (315 Hz and 395 Hz) of stimuli used in Experiment 1. ***B***, Comparison of averaged spectral amplitude of FFR at the stimulus frequencies among the conditions. Multiple comparisons revealed significantly greater responses for the Dich_Mono condition. This result can be explained by the difference in neural adaptation at the brainstem level due to the higher presentation rate in the Dich_TargetL/R conditions or Passive conditions compared with the Dich_Mono condition.

FFR amplitudes varied between left- and right-ear sounds and between some conditions related to the physical conditions of the stimuli (described later). What is critical for the scope of the current study is the difference between the Dich_TargetL and Dich_TargetR conditions within a left- or right-sound case. There was no significant difference between the Dich_TargetL and Dich_TargetR conditions in either right- or left-ear FFRs (Fig. 8B). This result is consistent with the previous studies which found no significant effect of attention on the averaged FFR (Galbraith, 1994; Varghese et al., 2015).

Less critically, we found that FFR amplitude was largest when the sound was presented monaurally (Mono_TargetL and Mono_TargetR conditions) compared with the binaural presentation conditions (Dich_TargetL, Dich_TargetR, and Dich_Passive conditions) (Fig. 8B). In the Mono conditions, the stimulus presentation rate (collective of the two ears) was lower than in the Dich conditions, which may induce less neural adaptation (thus, greater activity) in the auditory nuclei integrating the binaural information, e.g., in the inferior colliculus (IC), which is the main source of FFR (Chandrasekaran and Kraus, 2010; Bidelman, 2018).

Generally, FFR was larger for right-ear sounds than for left-ear sounds. This difference can be explained by the asymmetry in FFR amplitude between left and right ear sound presentations (Ballachanda et al., 1994) or the frequency dependency in FFR amplitude (Hoormann et al., 1992; Tichko & Skoe, 2017).

### FFR strength varied systematically around the MS onset

Next, we examined the relationship between the moment-by-moment FFR strength and the MS direction. This analysis was driven by our expectation that the MS direction may reflect the state of auditory attention at the MS occurrence, as suggested by the earlier sections. In this analysis, for a given MS towards the direction of interest, standard stimuli (separately for the left- and right-ear sounds) around the timing of the MS were found, and they were represented as point data expressing the FFR amplitude at the time relative to the MS. A moving Gaussian window (s = 50 ms, duration = 400 ms) along the time axis was used to average the pooled data across the MS occurrences, to obtain a smooth function representing a peri-MS FFR function, i.e., the time course of FFR amplitude around the MS (Fig. 9A, see Methods). Our main interest was the difference in the peri-MS FFR functions between when the MS direction was congruent and incongruent with the stimulus ear for which FFR was evaluated (referred to as the congruent and incongruent cases, respectively). Figures 9B and 9C show the normalized peri-MS FFR functions for the congruent and incongruent cases. Here, we summarized the data from all the participants (after excluding a few participants with poor signal-to-noise ratio) and for the Dich_TargetL and Dich_TargetR conditions. The normalization was performed to remove the effects of differences in FFR amplitude (in nV) that depended on participant and stimulus frequency (see Methods for details).

**Figure 9.**
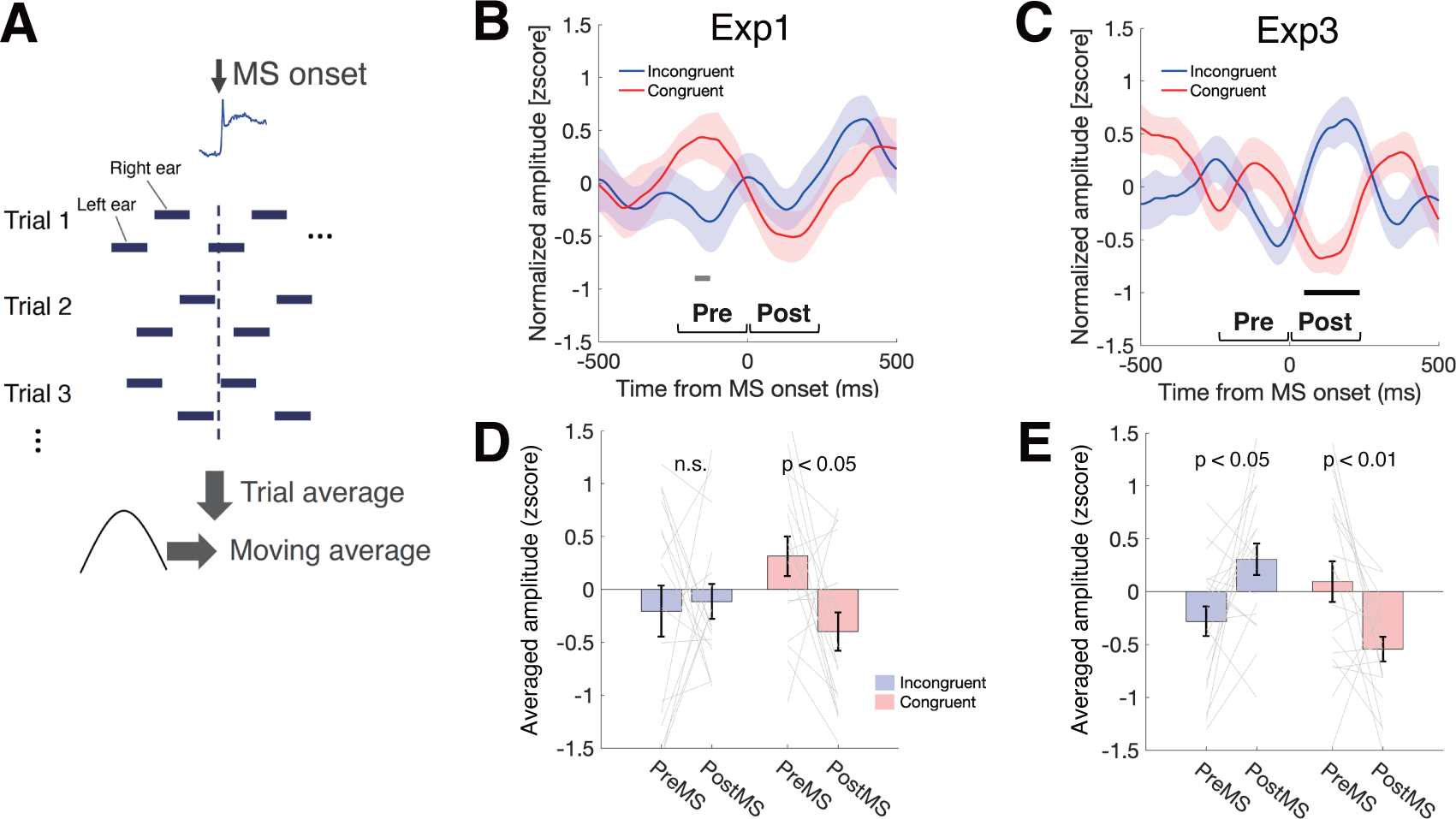
FFR changes time-locked to the MS onset. ***A***, Procedure for calculating peri-MS FFR changes. First, the components corresponding to the stimulus frequency of Fourier-transformed values of EEG data were extracted according to the MS onset. The extracted data were combined across trials while preserving the timing information (i.e., the timing of sound presentation around the MS). Then, the combined data were averaged with a moving Gaussian window. ***B***, The time course of peri-MS FFR changes for Experiment 1. Red and blue lines represent congruent and incongruent cases, respectively (e.g., the congruent case indicates the trials with rightward MS and the right-ear sound presentation). The black horizontal line indicates a significant cluster derived by the cluster-based permutation test between congruent and incongruent cases. ***C***, Comparisons of the FFR between pre- and post-MS for congruent (red bars) and incongruent cases (blue bars) for Experiment 1. ***D***, Same as ***B***, but for Experiment 3. ***E***, Same as ***C***, but for Experiment 3. The results indicate that FFR strength tended to decrease immediately after the MS occurrence for the congruent case.

We found that clusters from t = -168 ms to -135 ms for the normalized time courses were significantly larger for congruent than for incongruent cases in Experiment 1 (solid black lines in Fig. 9B). We calculated the change in FFR strength between pre-MS time range (-200 to 0 ms) and post-MS time range (0 to 200 ms). For experiment 1, we found no significant differences between pre-MS and pos-MS FFR for the incongruent case (p = 0.779, paired t-test, blue bars in Fig. 9C), while we found significant differences between pre-MS and pos-MS FFR for the congruent case (p = 0.0223, paired t-test, red bars in Fig. 9C). For Experiment 3, we found clearer dissociation of FFR changes after MS onset between congruent and incongruent cases (Fig. 9D). Clusters from t = 57 ms to 226 ms for the normalized time courses were significantly larger for incongruent than for congruent cases in Experiment 3 (solid black lines in Fig. 9D). There were significant differences between pre-MS and post-MS FFR strength for both congruent and incongruent cases (p = 0.0079, p = 0.0161, respectively, Fig. 9E). Overall, we found that FFR strength tended to increase and decrease immediately before and after MS onsets, respectively for the congruent case, and vice versa for the incongruent case. This result suggests that subcortical processes in the auditory system covary with the instantaneous fluctuation of attention states probably linked with microsaccade direction, which may not be captured by a whole trial average in the previous studies (Galbraith and Kane, 1993; Varghese et al., 2015).

We believe it unlikely that the MS-FFR relationship observed above was a byproduct of gaze shift during an auditory attention task via the MS’s correlation with gaze position in a compensatory manner (Fig. 5). There was no clear correlation between FFR amplitude and the gaze position when we divided the FFR data into the left-sided and right-sided gaze trials.

### The signal-to-noise ratio of FFR was highest for intermediate pupil size

We found a relationship between MS and FFR, suggesting that the MS and FFR were covaried by the auditory selective attention mechanisms. Did the arousal state, indicated by the pupil size, contribute to the FFR strength? An earlier study (McGinley et al., 2015) suggested that the intermediate pupil size reflects the optimal arousal state and is associated with neural responsiveness in the auditory cortex. In this analysis, the FFR data were divided into three classes of periods when the pupil size was small, mid, and large, with boundaries at 33.3 and 66.6 percentiles for each condition and participant (an example of boundaries for one participant is shown in the top panels of Fig. 10A. Two red lines represent the 33.3 and 66.6 percentiles, respectively). The pupil size classes were determined for each sound trial (180 ms; the period during which the left and right sounds were presented once, respectively). We found an association between the noise level in the FFR signals and the pupil size. Each panel in Fig.10B represents the spectral amplitude of the FFT signals during stimulus presentation (colored thick lines) and the noise floor (thick black lines calculated for the period -40 to 0 ms relative to the onset of standard sound). We note that the FFT signals during stimulus presentation were generally comparable across the pupil sizes, but the noise level tended to be smallest for the intermediate pupil size. Focusing on the amplitude at the stimulus frequency (between 315 and 395 Hz) confirmed this tendency (Fig. 10A).

**Figure 10.**
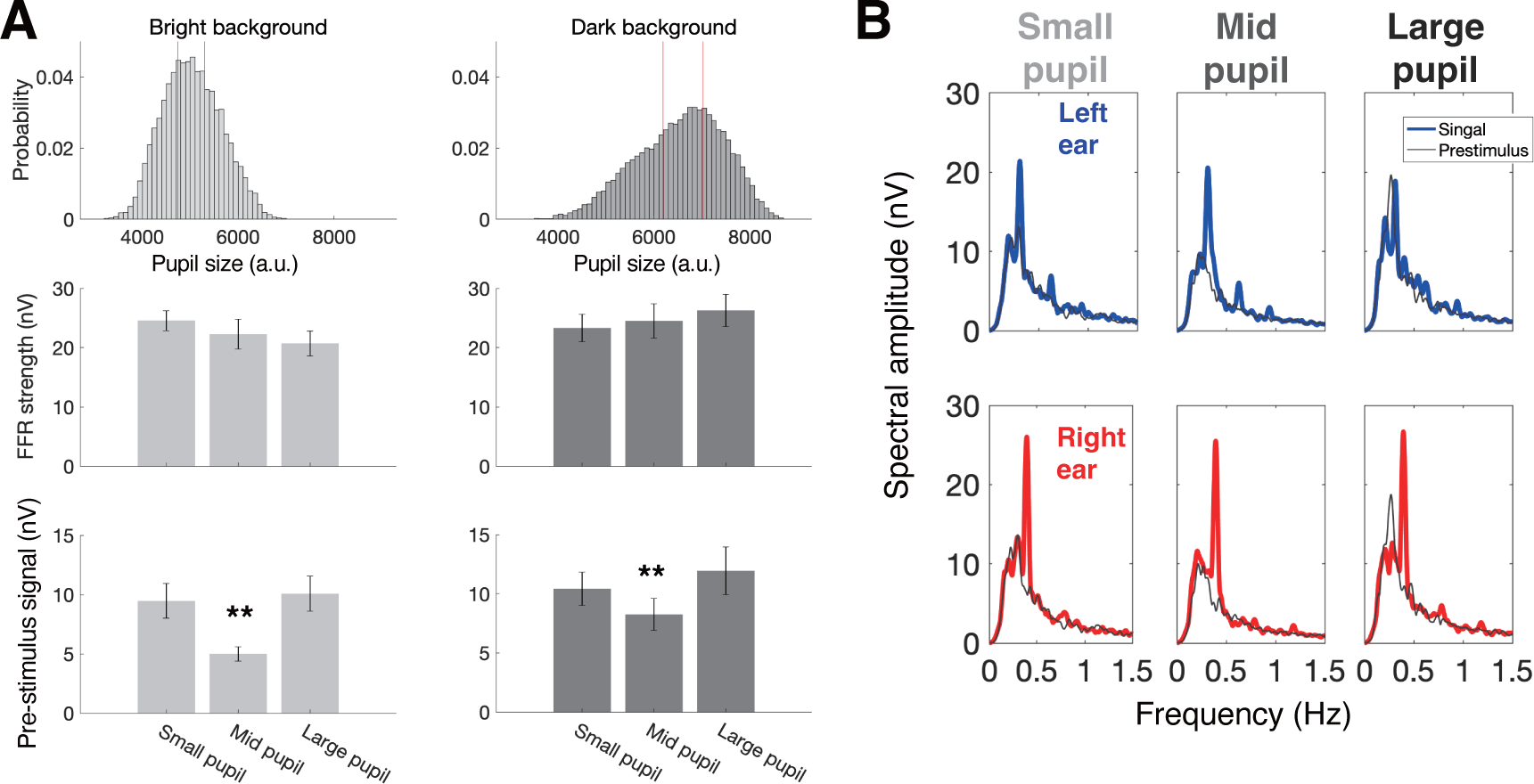
Relationship between FFR and pupil size. ***A***, Comparisons of FFR signal and pre-stimulus activity between three classes of pupil size (small, mid, and large). The top panels show the distributions of baseline pupil size for bright and dark display conditions, indicating that the pupil size was larger for the dark than for the bright conditions due to the pupillary light response. We defined the boundaries for three classes as the 33.3 and 66.6 percentiles of pupil size for each condition and each participant. The center panel shows the averaged FFR amplitudes for small, mid, and large pupil size classes. The bottom panel shows the averaged pre-stimulus activity, corresponding to the data point on the gray line at the stimulus frequency where the FFR exhibited the peak value (315 Hz for left-ear sound and 395 Hz for right-ear sound) in panel ***B***. Independently of the brightness of the display, pre-stimulus activity was significantly smaller for mid-pupil size compared with small and large pupil sizes. ***B***, The spectral amplitude of FFR for three classes of pupil size (small, mid, and large pupil size). Blue and red lines indicate the FFR to the left-ear and right-ear sounds, respectively. The gray lines indicate the pre-stimulus activity.

Taking the background luminance factor into account, two-way ANOVAs were conducted for FFR strength (center panels in Fig. 10B) and the noise floor (prestimulus activity, bottom panels in Fig. 10B). The factors were the pupil size (small, mid, and large) and background luminance (bright vs. dark). The data from 6 participants whose S/N ratio of averaged FFR was less than 0 dB for at least one of the following conditions were excluded from the statistical analysis; left-ear sound in the bright background, right-ear sound in the bright background, left-ear sound in the dark background, or right-ear sound in the dark background. First, we found no significant effects of pupil size or background luminance for FFR strength (ps > 0.1, center panels in Fig. 10B). Second, we found significant effects of pupil size (p = 0.0093, F(2,24) = 5.7140) and background luminance (p = 0.0180, F(1,12) = 7.4914) for the noise floor (bottom panels in Fig. 10B). The effect of background luminance indicated that the prestimulus subcortical activity was greater when the background luminance was darker. The effect of pupil size indicated that the noise level at the subcortical stage was the smallest for mid-pupil size. Thus, this result suggests that a pupil-linked optimal neural state also exists at the subcortical level, as found in the auditory cortex of mice (McGinley et al., 2015).

## Discussion

We found that the direction of microsaccade (MS) was generally biased contralateral to the ear to which the oddball sound was presented or that to which sustained auditory attention was directed. The stimulus-related modulation of MS bias after the oddball presentation was associated with performance in the auditory dichotic detection task. MS bias away from the location of sustained auditory attention included a component that compensates for the gaze shift. This compensation effect, however, cannot explain the relationship between stimulus-related MS bias and task performance. We also found that there was a relationship between subcortical frequency-following response (FFR) and MS. The results suggest that the auditory neural activity, even in the subcortical stage, fluctuates over time, probably with the states of attention/oculomotor system.

The microstructure of the MS time course after the presentation of auditory oddballs showed an early bias toward the stimulus and a late bias away from the stimulus. The early bias was only observed for the Dich_TargetBoth condition. Comparable characteristics of MS time course had been reported for the peripheral *visual* cue: MS direction tended to be biased toward the cued location in an earlier time range and be biased away from the cued location in a later time range for both humans and monkeys (Hafed et al., 2021; Hafed & Ignashchenkova, 2013; Laubrock et al., 2005; Tian et al., 2016; Tse et al., 2004). The early and late MS biases may be mediated by different neural circuits, involving the superior colliculus (SC) and frontal eye field (FEF), respectively: The inactivation of the monkey SC diminished the early MS bias toward the peripheral cue while the late opposite bias remained (Hafed & Ignashchenkova, 2013; Hafed et al., 2021). On the other hand, the inactivation of the monkey FEF influenced the late MS bias away from the cue, while the early MS bias toward the stimulus remained (Peel et al., 2016, Hafed et al., 2021). Our results extended this theory to auditory targets. It is important to note that the latencies of the early and late biases appear to be longer for the auditory target than for the visual target: The present results showed that the latencies in auditory-evoked MS bias were around 0.5 s and 1 s for the early and late biases, respectively, whereas the visually-evoked MS biases occurred around 0.2 s and 0.4 s, respectively. The latency difference between visually- and auditory-evoked MS bias suggests that the auditory-evoked MS bias may involve an indirect pathway from the auditory signal input to an oculomotor command, for example, through the connection between IC and SC, or the auditory input to the FEF (Kirchner et al., 2009).

Why was no early MS bias toward the oddball sounds observed for the conditions other than the Dich_TargetBoth condition, whereas the previous visual studies generally reported MS toward the visual cue (we call such an early MS bias “orienting MS”)? In the dichotic selective detection task, participants might employ a strategy inhibiting any stimulus-driven orienting response to reduce the probability of attentional capture by the oddballs on the unattended side. Few MSs toward the oddball sound in the Dich_TargetL/R conditions may reflect “success” in inhibiting unnecessary orienting responses toward the peripheral stimulus. The MS bias associated with the inhibition of orienting responses may be observable only for a task that requires sustained attention to prolonged stimuli (such as in the present study), and not for a task with transient cueing and stimuli (as in the earlier studies described above).

Our findings on the link between the MS bias and task performance are consistent with the idea that the early MS bias is suppressed during a task that requires sustained spatial selective attention. The MS tended to be biased toward the position of auditory oddballs for the low-performance trials (false alarm and long RT trials), suggesting that the suppression of the orienting MS failed when the participants’ alertness level was low. Such an inhibition of orienting responses might be controlled by the top-down signal from FEF (Van der Stigchel et al., 2012). The result implies that the MS can be an index of time-varying attention states relevant to spatial processing in the auditory domain.

The current study showed that the gaze-compensatory MS was accompanied by spontaneous gaze bias towards the attended direction during a dichotic selective detection task. One might argue that the link between MS and participants’ task performance can be explained by the gaze shift. Indeed, an earlier study reported the gaze shift depended on spatial attention for both the visual and auditory domains (Gopher, 1973). Previous neurophysiological studies showed that the eye gaze position influences the activity of the inferior colliculus and the primary auditory cortex in monkeys (Groh et al., 2001; Werner-Reiss et al., 2003; Fu et al., 2004; Bulkin and Groh, 2012). In humans, previous studies found that directing the gaze toward a sound position enhances discrimination of auditory spatial information or deviant detection in a target auditory stream (Maddox et al., 2014; Pomper and Chait, 2017). Although the participants were instructed to fixate on the center of the display, they might implicitly take advantage of the spontaneous gaze shift to enhance task-related auditory processing. However, the factor influencing MS was not only the gaze position but also the presentation of the oddball sound, as discussed above. More critically, the effect of RT was not significant for the gaze position around the oddball sound (Fig.6C), while it was significant for the MS time course (Fig.4C). Therefore, the present findings on stimulus-related MS bias cannot be explained solely by gaze-compensatory MS.

Finally, we asked whether the MS can be an index of fluctuating neural activity which may co-vary with the attention state. We measured the auditory subcortical responses (FFRs) during a dichotic selective detection task. When the data were averaged over the entire period of stimulus presentation, there was no significant effect of attentional set on the FFR to standard sounds for both left- and right-ear sounds. This result is consistent with the null effect of attention on FFR reported by previous studies (Galbraith and Kane, 1993; Varghese et al., 2015). Through careful inspection, however, we found that the FFR at the stimulus frequency was linked with the onset and direction of MS during a selective detection task. FFR may be susceptible to temporal variation in the attention states (Esterman et al., 2013; Terashima et al., 2021; van den Brink et al., 2016), which co-varies with the eye activities as revealed by the current study. The current result may shed light on the discrepant results of studies about the attention effect on FFR. The temporal fluctuation of the subcortical activity may co-vary with the attention states, which cannot be captured by a simple whole trial average in conventional FFR studies but might be indexed by the MS. Recent studies found the effect of attention on FFR signal to a running speech stimulus (Etard et al., 2019; Forte et al., 2017) and by applying source-level analysis of EEG data to emphasize the subcortical activity (Price and Bidelman, 2021), which may be able to capture the time-varying fluctuation of the effect of attention. The result showing the link between FFR and pupil size also supports the idea that neural activities in the subcortical stage of the auditory pathway fluctuate depending on the states of attention or arousal (Fig. 10).

## Author contributions

S.Y. and S.F. designed research; S.Y. performed experiments; S.Y. analyzed data; and S.Y. and S.F. wrote the paper.

